# Phylodynamics Uncovers the Transmission of Antibiotic-Resistant *Escherichia coli* between Canines and Humans in an Urban Environment

**DOI:** 10.1101/2023.06.01.543064

**Authors:** Nikolina Walas, Nicola F. Müller, Emily Parker, Abigail Henderson, Drew Capone, Joe Brown, Troy Barker, Jay P. Graham

**Author notes:** Correspondence: Nikolina Walas.

## Abstract

The role of canines in transmitting antibiotic resistant bacteria to humans in the urban environment is poorly understood. To elucidate this role, we utilized genomic sequencing and phylogenetics to characterize the burden and transmission dynamics of antibiotic resistant *Escherichia coli* (ABR-Ec) cultured from canine and human feces present on urban sidewalks in San Francisco, California. We collected a total of fifty-nine ABR-Ec from human (n=12) and canine (n=47) fecal samples from the Tenderloin and South of Market (SoMa) neighborhoods of San Francisco. We then analyzed phenotypic and genotypic antibiotic resistance (ABR) of the isolates, as well as clonal relationships based on cgMLST and single nucleotide polymorphisms (SNPs) of the core genomes. Using Bayesian inference, we reconstructed the transmission dynamics between humans and canines from multiple local outbreak clusters using the marginal structured coalescent approximation (MASCOT). Overall, we found human and canine samples to carry similar amounts and profiles of ABR genes. Our results provide evidence for multiple transmission events of ABR-Ec between humans and canines. In particular, we found one instance of likely transmission from canines to humans as well as an additional local outbreak cluster consisting of one canine and one human sample. Based on this analysis, it appears that canine feces act as an important reservoir of clinically relevant ABR-Ec within the urban environment. Our findings support that public health measures should continue to emphasize proper canine feces disposal practices, access to public toilets and sidewalk and street cleaning. *Importance:* Antibiotic resistance in *E. coli* is a growing public health concern with global attributable deaths projected to reach millions annually. Current research has focused heavily on clinical routes of antibiotic resistance transmission to design interventions while the role of alternative reservoirs such as domesticated animals remain less well understood. Our results suggest canines are part of the transmission network that disseminates high-risk multidrug resistance in *E. coli* within the urban San Francisco community. As such, this study highlights the need to consider canines, and potentially domesticated animals more broadly, when designing interventions to reduce the prevalence of antibiotic resistance in the community. Additionally, it showcases the utility of genomic epidemiology to reconstruct the pathways by which antimicrobial resistance spreads.

## Introduction

Antibiotic resistance (ABR) is a global health crisis with more than 28 million antibiotic-resistant infections occurring in the US each year (1). In 2019, *E. coli* was estimated to be the top contributor for deaths attributable to bacterial ABR (2). Despite the increasing rates of ABR bacterial infections, these pathogenic species targeted by antibiotics constitute a small proportion of the gut microbiome. Nonetheless, resistance to common antibiotics can pass between this population and the normal gut flora or surrounding environmental bacteria (3). Studies have shown *E. coli,* a key species associated with ABR, to be a member of the gut microbiome in about 90% of individuals (4), (5). Rates of resistance in *E. coli* have been reported to range from 8.4% to 92.9% and models predict over half of *E. coli* invasive species may become 3^rd^ generation cephalosporin resistant by 2030 (6). Increasing rates of resistance in such populations have been attributed to circulation of specific resistant bacterial clonal species as well as transmission of antibiotic resistance genes (ARGs), often mediated by mobile genetic elements such as integrons, transposons and plasmids (7), (8).

The global dissemination of antibiotic resistant bacteria (ARB) such as *E. coli* is best described using the One-Health paradigm where human, animals and the environment act as overlapping pillars of transmission (8), (9). Despite this, research has focused heavily on clinical interventions to reduce ABR transmission in human pathogens. Studies have suggested animals and the environment play equally important roles as reservoirs (10), but the extent of their contribution to the ARB transmission cycle is still not well understood (9). It is known, however, that runoff contaminated with ARGs from human and livestock waste exacerbates the natural exchange of resistance genes from environmental bacteria to animal and human pathogens (11).

Domesticated animals have long been established as a reservoir for ARB but there is still controversy regarding the exact role canines play in transmission to humans and environmental contamination. The species has been shown to act as a vector for clinically relevant extended spectrum β-lactamases (ESBL) in urban settings (12). With regards to humans, various cohabitation studies have shown a wide range of overlap in the resistance profile of domesticated canines and their respective owners (13), (14). For example, Johnson et al. showed that a strain of uropathogenic *E. coli* appeared to move between a canine and humans in the same household (15). These findings have not been reproducible in different settings, most likely due to limitations in sampling time and the transient nature of the gut microbiome (16).

Another challenge in studying the burden of animal species on ARB transmission to humans is the lack of computational tools to robustly infer transmission directionality. The majority of studies investigating the role of animals use the co-occurrence of resistance as an indicator of spread or rely on methods that are insufficient to conclude directionality (17). For example, vertical evolution of shared bacterial species has traditionally been investigated using phylogenetic modeling. However, these analyses are usually constrained to the core genome of bacteria due to the high degree of conservation (18) and largely ignore metadata during model generation (19). Phylodynamics, which uses genetic data to infer epidemiological dynamics, is an increasingly popular statistical framework that can be used to infer transmission events. It has remained largely overlooked and underdeveloped in its application to bacterial pathogens (20) due to their often more complex evolutionary mechanisms, compared to, for example, viral pathogens (19). Understanding the transmission dynamics of ABR-Ec between species will help to inform disease control strategies and public health interventions.

Most studies investigating the role of animals in ARB have been conducted in low-and middle-income countries (LMIC) that are known to bear the greatest burden of ABR due to a range of political, economic, and infrastructure factors (9), (21). Few studies (12) have detailed the resistance profiles of canines with regards to its contribution to environmental contamination of ARGs and risk of transmission to humans in high-income (HIC) urban settings. Therefore, the aim of this study was to characterize the prevalence of ABR-Ec in human and putative canine fecal samples found on the sidewalks of San Francisco, CA, USA where the population of unhoused individuals is relatively high. Results from this study can be used to better understand the public health risk from canine fecal contamination and provide evidence of transmission directionality between canines and humans using a phylodynamic approach.

## Materials and Methods

### Sample Collection and DNA Isolation

Fecal samples were located based on open defecation hotspots that were determined by San Francisco’s 311 municipal system that allows citizens to report issues to the city’s department of public works. Samples were collected from a perimeter of 20 blocks in the Tenderloin and SOMA neighborhoods and included sidewalks on either side of the street. Biospecimens were collected on Wednesday mornings before street cleaning in September and October of 2020. Samples were placed into one-liter Whirl-Pak® Sample Bags (Millipore Sigma, Darmstadt, Germany) and stored in a cooler with ice packs for a maximum of 4 hours before being transferred and stored in 1.5 mL cryotubes at −20 °C.

The QIAamp 96 Virus QIAcube HT Kit was used to purify DNA. DNA was first extracted from 100 mg of stool using a bead beating tube and 1 mL Qiagen Buffer ASL. As a positive control for DNA extraction, G-block from IDT matching sequence for phocine herpesvirus was spiked in. The bead beating tubes were vortexed for 5 minutes, incubated at room temperature for 15 minutes, and centrifuged at 14,000 rpm for 2 minutes, in alignment with previous study methods (22). Extraction was completed using the QIAamp 96 Virus 281 QIAcube HT Kit and done on the QIAcube. The extracted nucleic acids were stored at 4 °C for less than 24 hours before being stored at −80 °C.

Human fecal samples were determined using a previously validated dPCR method to detect human mitochondrial DNA (mtDNA) (23). A QIAGEN QIAcuity Four machine was used to run 40 µl reactions on QIAcuity Nanoplate 26K 24-wll plates. Reactions included 2 µl template, 27.2 µl nucleotide-free H20, 1X CIAcuity PCR MasterMix, 400 nM sun probe, and 800 nM of forward and reverse primers. Cycles consisted of 2 minutes at 95 °C, 40 cycles of 15 seconds at 95 °C, and 30 seconds at 59 °C. Positive and negative controls were run each day of analysis. Quibit dsDNA results were used to normalize gene copy estimates with a positive cut-off log-adjusted value of 1.

### Phenotypic Analysis

In order to identify resistant isolates, fecal samples were first streaked on three plates containing MacConkey, MacConkey and Ampicillin (32 µg/ml), or MacConkey and Ceftriaxone (1 µg/ml) mediums. Putative *E. coli* isolates determined by indole testing were selected from each plate for susceptibility testing. The antibiotic resistance pattern of the isolates was determined using a disc diffusion panel of ten antibiotics in accordance with the Clinical and Laboratory Standards Institute (CLSI) guidelines (24): ampicillin (10µg disc), ertapenem (10µg disc), ciprofloxacin (5 µg disc), trimethoprim-sulfamethoxazole (25µg, disc), nitrofurantoin (300µg, disc), cefepime (30µg, disc), piperacillin-tazobactam (110µg, disc), cefazolin (30µg, disc), cefotaxime (30µg, disc) and ceftazidime (30µg, disc). Isolates resistant to at least one antibiotic and with a unique resistance profile were selected for whole genome sequencing.

### Whole Genome Sequencing, Assembly, and Analysis

DNA for whole genome sequencing was purified from ABR isolates using the Qiagen DNeasy Blood and Tissue extraction kit. Whole-genome sequencing data was generated using an Illumina NovaSeq 6000 platform with a paired-end protocol (Nextera XT library; Illumina). Quality of reads was assessed using FastQC v0.11. De novo assemblies of the paired short reads were generated using Unicycler v.0.3.0b which acts as a SPAdes optimizer (25). Contigs below 500 bp were excluded from the final draft assemblies. Quality of assembled sequences was assessed using QUAST v5.0 (26).

Antibiotic resistance genes, plasmid types, and virulence genes were identified using the ABRicate tool (version 1.0) (27). The ResFinder database was used to detect resistance genes with a 90% minimum match and 80% minimum length. The PlasmidFinder database was used to detect the plasmid replicons in isolates with an 80% minimum coverage and identity. The VFDB and Ecoli_VF databases were used to detect virulence genes with an 80% minimum coverage and identity. Pathotypes of diarrheagenic *E. coli* were screened according to the presence of previously defined virulence genes (28). Plasmid content of the isolates was determined by assigning contigs of draft genomes as plasmid or chromosomal using mlplasmids (29) and MOB-suite (30). Contigs smaller than 1000 base pairs were filtered out of the analysis. The default parameter for *E. coli* plasmids was used for MOB-suite analysis. MOB-suite results were used in place of mlplasmids for contig calls with a minimum posterior probability below 75% from mlplasmids. Contigs with discrepant results from the two programs were not included in the final analysis.

MLST version 2.19 (31) was used to perform *in silico* multilocus sequence typing (MLST), based on seven housekeeping genes (adk, fumC, gyrB, icd, mdh, purA, and recA). cgMLSTFinder version 1.1 (32), (33) accessed through the Center for Genomic Epidemiology was used to assign the cgMLST to each *E. coli* isolate. Isolates were considered epidemiologically linked with a genetic distance, calculated by the number of allele differences divided by the number of alleles shared between two isolates, below 0.0105 according to previous methods (34). Phylogroup assignment was determined using the *in silico* Clermont 2013 PCR typing method tool EzClermont (35).

### Phylogenetic and MASCOT Analysis

Assembled draft genomes were annotated using Prokka version 1.12 (36). Pan-genome analysis was completed using ROARY version 3.13 (37) with a core gene defined as being present in over 99% of the isolates. Core genome alignment comprised of 2905 core genes was generated using the MAFFT setting in ROARY. SNP-sites (38) was used to extract 200,473 single-nucleotide polymorphisms (SNPs) from the core genome alignment. A maximum-likelihood tree was then calculated using RAxML version 8.2.12 (39) with the general time-reversible model (GTRCAT) and 100 bootstrap replicates. Recombination events detected by ClonalFrameML version 1.12 (40) were masked to produce a recombination free phylogenetic tree. The tree was then visualized in iTol version 6.5.8 (41).

To infer the transmission dynamics between canines and humans, we first split the dataset into local outbreak clusters according to previous methods (41), (42). We defined a local outbreak cluster as any set of sequences that are at most 200 SNPs apart. MASCOT was then used jointly on all outbreak clusters inferring the effective population size of *E. coli* in humans and canines, the rates of transmission between them, and the rate of introduction of *E. coli* into either compartment. We assumed a rate of evolution of 4 SNP’s per year, based on the rate estimate for *Shigella sonnei* from previous literature (44). As a site model, we used a GTR+Γ_4_ with estimated rates. Additionally, we reconstructed the host type of internal nodes in the local outbreak clusters, as well as the posterior distribution of host jumps, as the number of edges for which parent and child node are inferred to be in different hosts. Tree output was visualized using densitree (45) and ggtree (46).

### Statistical Analysis and Data Visualization

The quantitative output of the number of ARGs in a species was transformed and treated as categorical variables in the statistical tests. Contingency tables based on ARG presence or absence in the species were generated and analyzed using the non-parametric two-tailed Fischer’s Exact test due to the small sample size. The same methodology was used to analyze virulence gene presence. The significance value was set at 5%.

All data manipulation and analyses were done using R Studio Software version 1.4.1073 in tandem with the following R packages: ggplot (47), dplyr (48), stringr (49), tidyr (50), and ggsankey (51). Spatial distribution of sample sites was visualized using QGIS(52) with a pseudo-Mercator projection.

### Data Availability

The draft assembled genomes for this project have been deposited in a DDBJ/ENA/GenBank Bioproject under the accession PRJNA910158. The version described in this paper is version 1. Individual genome sequences can be found with the following link: https://www.ncbi.nlm.nih.gov/bioproject/PRJNA910158

## Results

### Canine and human fecal sample characteristics

We sampled fifty-nine fecal samples over the course of five weeks from September to October 2020 in San Francisco, CA, USA. Fecal samples were assumed to be human based on visual discrimination by field staff. The range of sample sites covered a distance of 1 mile and spanned from the Tenderloin district to south of Market Street (Figure 1). Despite selectively collecting human-like fecal samples, only 20% (12/59) of samples were human based on the mtDNA analysis (53), (23). Due to the large size and visual appearance, all non-human fecal samples were assumed to be putative canine feces, which were commonly observed in the area. About 25% (3/12) of human fecal samples did not contain any antibiotic resistant *E. coli* (ABR-Ec) compared to 48% (23/47) of canine fecal samples. More than one phenotypically unique ABR-Ec was isolated from 58% (7/12) of human fecal samples and 19% (9/47) of canine samples.

**Figure 1:**
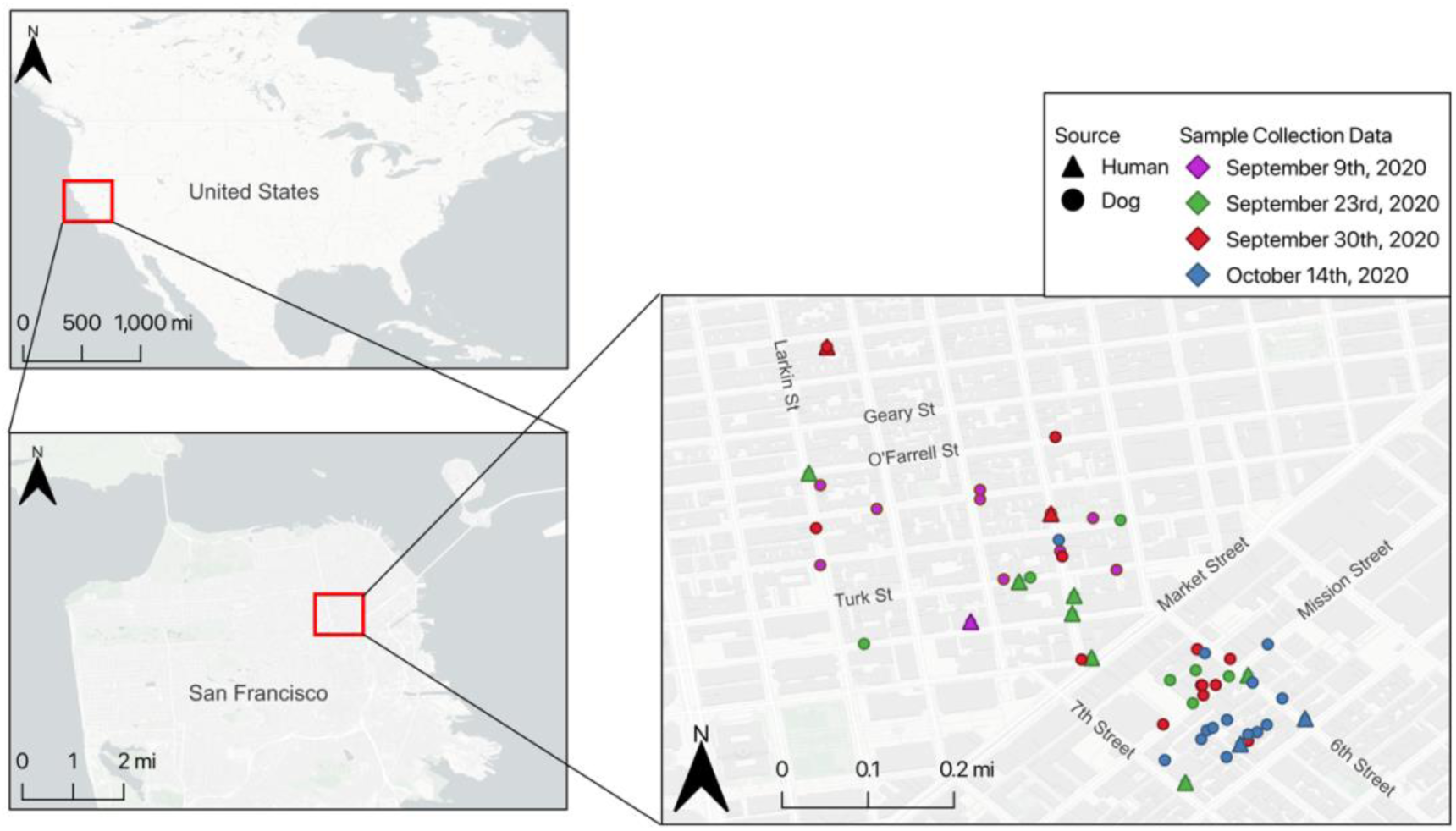
Fifty-nine fecal samples were collected over a one-mile range in the Tenderloin and SoMa neighborhoods of San Francisco, CA, USA. Samples were collected over one month period on four collection dates. Spatial distribution of sample sites was visualized using QGIS in a Pseudo-Mercator projection.

### Resistance, Plasmid Carriage, and Pathotypes in Human and Canines

The resistant isolates from both sources showed a similar distribution in phenotypic resistance (Figure 2A). Close to 100% of the isolates from both sources were phenotypically resistant to the penicillin, ampicillin, and the first-generation cephalosporin, cefazolin. A larger proportion of human isolates compared to canine isolates were phenotypically resistant to the trimethoprim-sulfamethoxazole (75%, 52%) and ciprofloxacin (40%, 8%) respectively.

**Figure 2:**
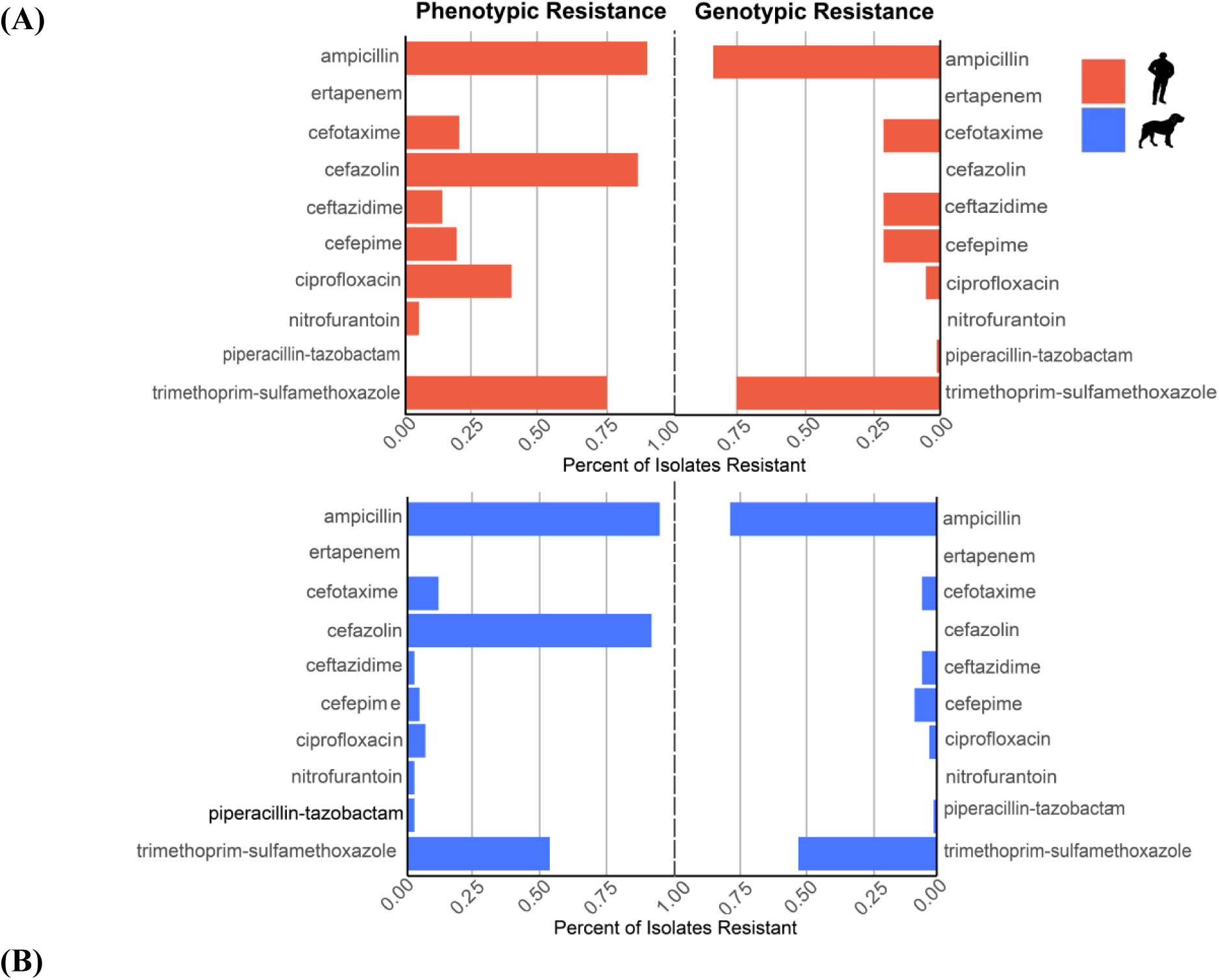
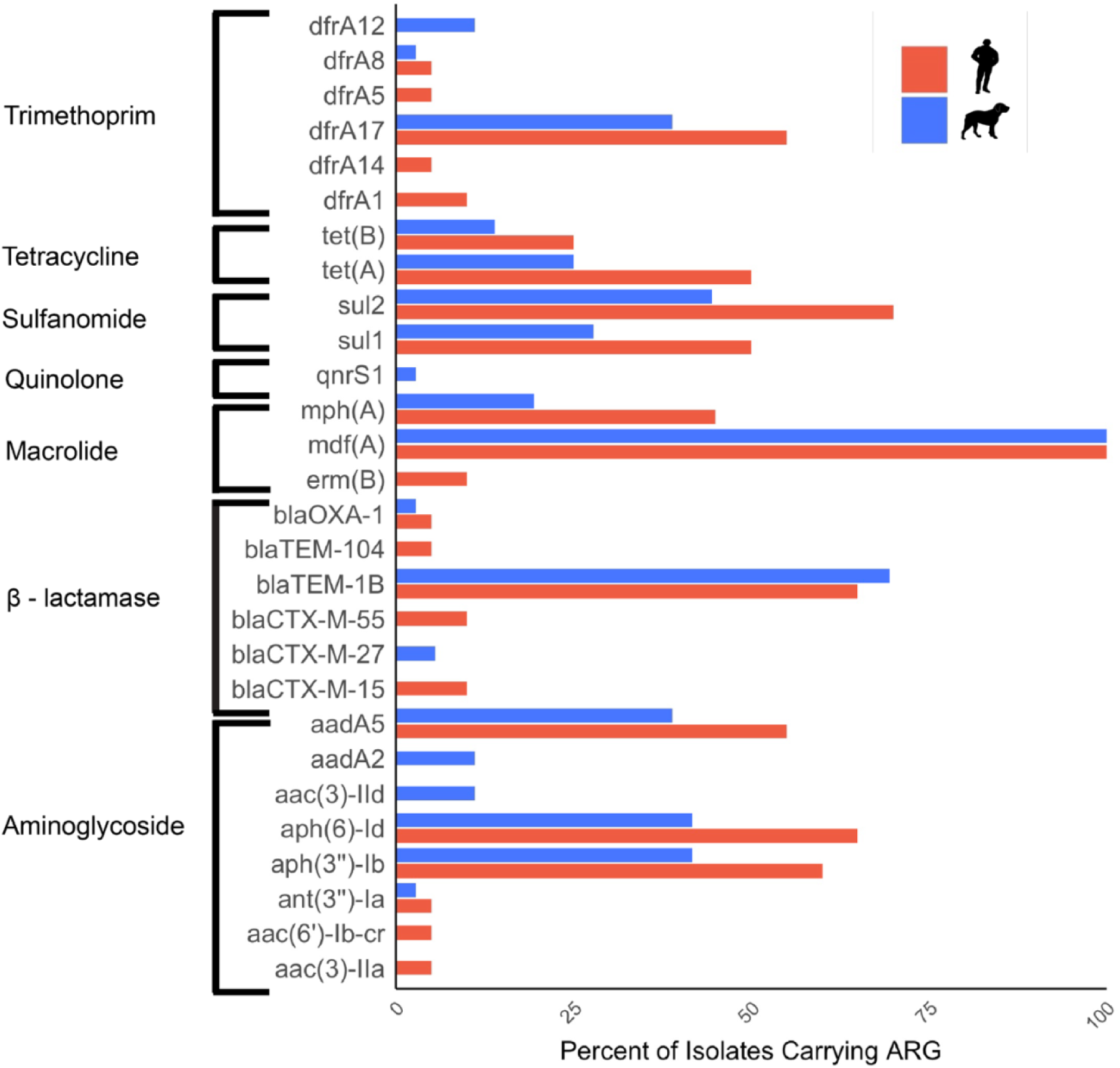
Isolates from humans and dogs were selectively plated on MacConkey, MacConkey and Ampicillin, or MacConkey and Ceftriaxone, and assessed for phenotypic resistance (A). Sequences of isolates determined by whole genome sequencing were analyzed using the ResFinder database with ABRicate (B).

Phenotypic resistance to the third and fourth generations of cephalosporin drugs cefotaxime, ceftazidime, and cefepime was higher in humans (20%, 20%, 15%) compared to canines (5%, 14%, 3%). Despite being phenotypically resistant to cefazolin in almost 100% of the isolates, none of the isolates carried cefazolin specific resistance genes.

With regards to antibiotic resistance genes (ARGs), we identified a total of 28 ARGs, with 5 ARGs found only in humans and 9 ARGs found only in canines. Although only 50% of ARGS were shared by both species, there was no significant difference in the presence of resistance genes between two species (Table 1). The mean number of ARGs per isolate was 7.20 in human and 5.11 in canines. The most common resistance gene was the broad spectrum *mdf(A)* gene, present in all isolates, followed by the β-lactamase gene *bla*_TEM-1B_ present in 65% (n=13) and 70% (n=25) of isolates in humans and canines, respectively. The class A β-lactamase *bla*_TEM104_ was only present in one human isolate. There was no overlap in the carriage of extended-spectrum β-lactamase (ESBL) genes between the two species (Figure 2B). The only ESBL gene detected in canine isolates was *bla*_CTX-M-27_ while human isolates carried *bla*_CTX-M-55_ and *bla*_CTX-M-15_.

**Table 1:** Distribution of ARGs between Human and Canine Isolates

| Resistance Group | Gene | Human (%) | Canine (%) | p-value* |
| --- | --- | --- | --- | --- |
| Aminoglycoside | *aac(3)-IIa* | 5.0 | 0.0 | 0.357 |
|  | *aac(6')-Ib-cr* | 5.0 | 0.0 | 1.000 |
|  | *ant(3'')-Ia* | 5.0 | 2.78 | 1.000 |
|  | *aph(3'')-Ib* | 60.0 | 41.7 | 0.266 |
|  | *aph(6)-Id* | 65.0 | 41.7 | 0.163 |
|  | *aac(3)-IId* | 0.0 | 11.1 | 0.285 |
|  | *aadA2* | 0.0 | 11.1 | 0.285 |
|  | *aadA5* | 55.0 | 38.9 | 0.428 |
| β-lactamase | *bla<sub>CTX-M-15</sub>* | 10.0 | 0.0 | 0.123 |
|  | *bla<sub>CTX-M-27</sub>* | 0.0 | 5.6 | 0.533 |
|  | *bla<sub>CTX-M-55</sub>* | 10.0 | 0.0 | 0.123 |
|  | *bla<sub>OXA-1</sub>* | 5.0 | 2.8 | 1.000 |
|  | *bla*<sub>TEM-1B</sub> | 65.0 | 69.4 | 0.772 |
|  | *bla*<sub>TEM-104</sub> | 5.0 | 0.0 | 0.357 |
| Macrolide | *erm*(B) | 10.0 | 0.0 | 0.123 |
|  | *mdf*(A) | 100.0 | 100.0 | 1.000 |
|  | *mph*(A) | 45.0 | 19.4 | 0.064 |
| Quinolone | *qnrS1* | 0.0 | 2.8 | 1.000 |
| Sulfonamide | *sul1* | 50.0 | 27.8 | 0.146 |
|  | *sul2* | 70.0 | 44.4 | 0.095 |
| Tetracycline | *tet*(A) | 50.0 | 25.0 | 0.080 |
|  | *tet*(B) | 25.0 | 13.9 | 0.468 |
| Trimethoprim | *dfrA1* | 10.0 | 0.0 | 0.123 |
|  | *dfrA5* | 5.0 | 0.0 | 0.357 |
|  | *dfrA8* | 5.0 | 2.8 | 1.000 |
|  | *dfrA12* | 0.0 | 11.1 | 0.285 |
|  | *dfrA14* | 5.0 | 0.0 | 0.357 |
|  | *dfrA17* | 55.0 | 38.9 | 0.275 |

\*alpha value of 0.05

We detected twenty-seven plasmid replicons, six of which were unique to canines and ten to humans (supplemental table 1). Col156 was the most prevalent replicon in human isolates (n=15) and second most prevalent in canine isolates (n=15). IncFIB was the most prevalent in canine isolates (n=20) and third most prevalent in human isolates (n=13). In terms of ARG localization, a greater proportion of ARGs in human isolates were plasmid bound compared to canine isolates (Figure 3). Only one of the detectable β-lactamase genes, *bla*_CTX-M-15_, in human isolates was chromosomal. Seven of *bla*_OXA-1_ (n=1) and *bla*_TEM-1B_ (n=6) β-lactamase genes were chromosomal in canines. The ESBL genes *bla*_CTX-M-27_ (n=1) and *bla*_CTX-M-55_ (n=2) in canine and human isolates, respectively, were plasmid-bound. Plasmid replicon type could not be identified for the majority of resistance genes carrying plasmid contigs. Human fecal samples carried an IncK2/Z plasmid contig carrying sulfonamide, trimethoprim, and aminoglycoside genes. Three of the seven plasmid replicons were shared between species. We found plasmid contigs carrying *bla*_TEM-1B_ and identified as IncFIA, IncFIA/IncFIC and IncFIA/IncFII in human (n=1, n=2, n=1) and canine (n=3, n=4, n=1) isolates (Table 2).

**Table 2:** Distribution of ARG Carrying Plasmid Replicons Between Human and Canine Isolates

| Plasmid Replicon Type | Resistance Genes | Number of Isolates (n) | Sequence Types (n) |
| --- | --- | --- | --- |
| IncFIA | *bla*<sub>TEM-1B</sub> | Human (1)<br>Canine (3) | 131 (1), 29 (1), 69 (2) |
| IncFIA/IncFIB/IncFIC | *bla*<sub>TEM-1B</sub> | Human (1) | 58 (1) |
| IncFIA/IncFIC | *bla*<sub>TEM-1B</sub> | Human (2)<br>Canine (3) | 29 (2), 569 (1), 10 (2) |
|  | *bla*<sub>TEM-1B</sub>, *aac*(3'')-IIb | Canine (1) | 162(1) |
| IncFIA/IncFII | *bla*<sub>TEM-1B</sub> | Canine (1) | 69 |
|  | *bla*<sub>TEM-1B</sub>, *aph*(3'')-Ib, *aph*(6)-Id, *sul2* | Human (1) | 10 |
| IncFIB/IncI-gamma/K1 | *bla*<sub>TEM-1B</sub> | Canine (1) | 8857 (1) |
| IncFIC | *bla*<sub>TEM-1B</sub> | Canine (2) | 969 (2) |
| IncK2/Z | *aph*(6)-Id, *dfrA14*, *sul2* | Human (1) | 131 (1) |

**Table 3:** Epidemiologically linked multidrug resistant (MDR) isolates

| Sharing Group | Number of isolates | ST | cgST | Avg allele difference | Phenotypic Resistance Profile | Genotypic Resistance Profile | Sample mtDNA (n) | Sampling Date |
| --- | --- | --- | --- | --- | --- | --- | --- | --- |
| 1 | 2 | 131 | 136431 | 0 | AMP, KZ, SXT | *aac(3)-IId_1, aadA2, bla<sub>TEM-1B</sub>, dfrA12, mdfA, mphA, sul1_5, tetA* | Canine | 9/30/20 |
| 2 | 2 | 69 | 133094 | 1 | AMP, KZ | *bla<sub>TEM-1B</sub>, mdfA* | Canine | 9/23/20<br>9/30/20 |
| 3 | 2 | 969 | 40914 | 0 | AMP, KZ | *bla<sub>TEM-1B</sub>, mdfA, tetA*<br>*bla<sub>TEM-1B</sub>, mdfA* | Canine | 9/23/20 |
| 4 | 2 | 155 | 62601 | 2 | AMP, KZ, SXT | *aadA2, aph(3'')-Ib, aph(6)-Id, bla<sub>TEM-1B</sub>, dfrA12, mdfA, sul1_5, sul2_3,* | Canine | 9/30/20 |
| 5 | 2 | 10 | 78278 | 1 | AMP, SXT | *aadA5, aph(3'')-Ib, aph(6)-Id, bla<sub>TEM-1B</sub>, dfrA17, mdfA, mphA, sul1_5, sul2_2, tetA,* | Canine | 10/14/20 |
| 6 | 6 | 2541 | 130412 | 2 | AMP, KZ, SXT<br>AMP, CTX, KZ,<br>SXT | *aadA5, aph(3'')-Ib, aph(6)-Id, bla<sub>TEM-1B</sub>, dfrA17, mdfA, sul2_14*<br>*aadA5, aph(3'')-Ib, aph(6)-Id, bla<sub>TEM-1B</sub>, dfrA17, mdfA, sul2_21* | Human (1)<br>Canine (5) | 9/23/20,<br>9/30/20,<br>10/14/20 |

Pre-defined virulence genes used to identify the six diarrheagenic *E. coli* pathotypes determined that one canine isolate belonging to ST131 carried the *daaE* gene indicative of diffuse adherent *E. coli*. About 25% of human and canine isolates carried the virulence gene *eae*. Virulence gene carriage was not significantly different between the two species (Supplemental table 2).

### Clonal and Phylogenetic Relationships Spanned Species

Twenty-eight sequence types were detected, including the well-known pandemic lineages 131, 69 and 1193 (Figure 4). Human isolates spanned 11 STs and canine isolates spanned 17 STs. The most common phylogroup was A (n=17) followed by B2 (n=15) and B1 (n=13).

**Figure 3:**
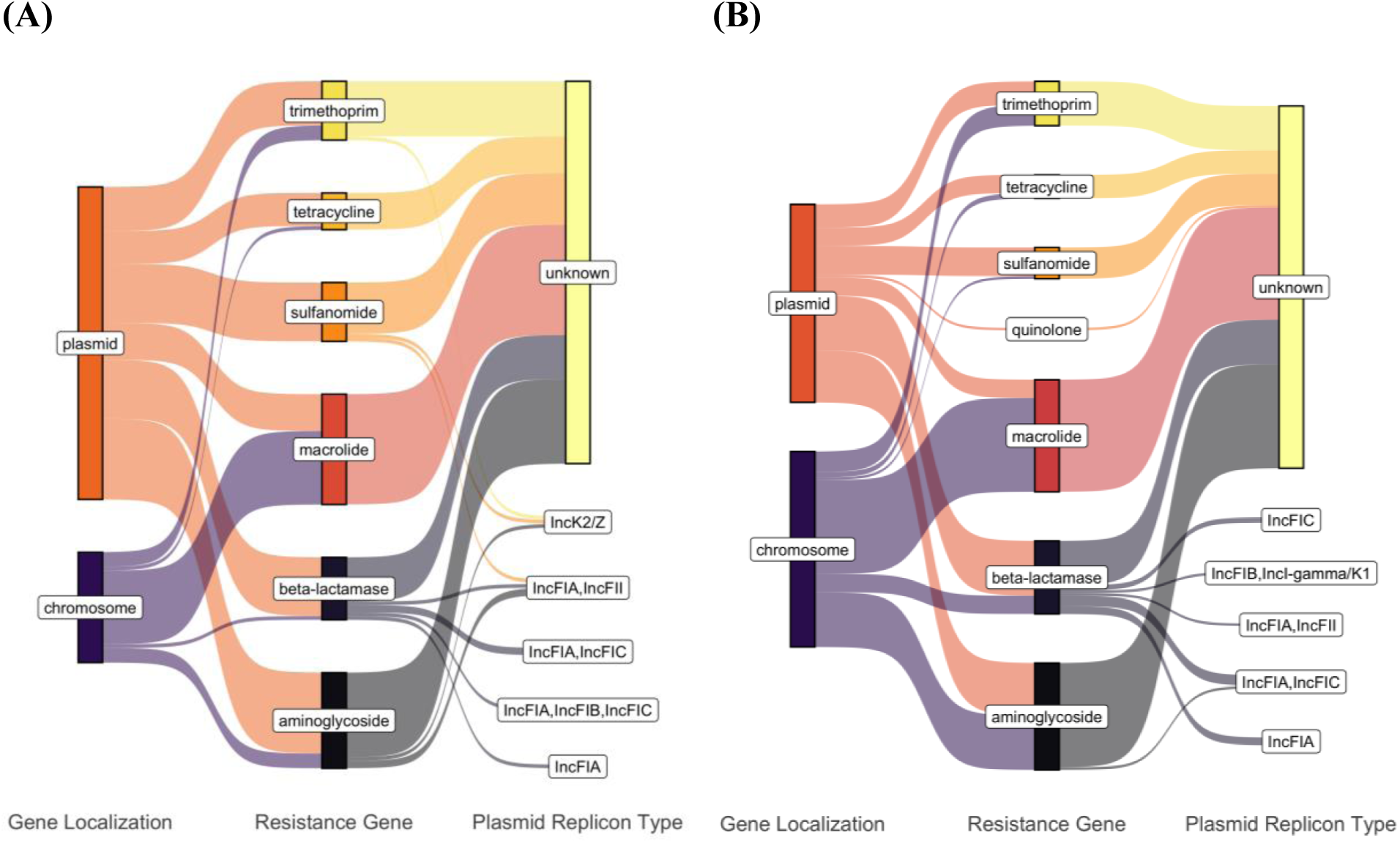
Contigs containing ARGs determined by ResFinder were assigned as plasmid or chromosomal according to mlplasmids and MOB-suite in human (A) and dog (B) isolates. Replicon types of the contigs were determined by MOB-suite. Distribution of localization and replicon types was visualized using ggsankey in R studio.

**Figure 4:**
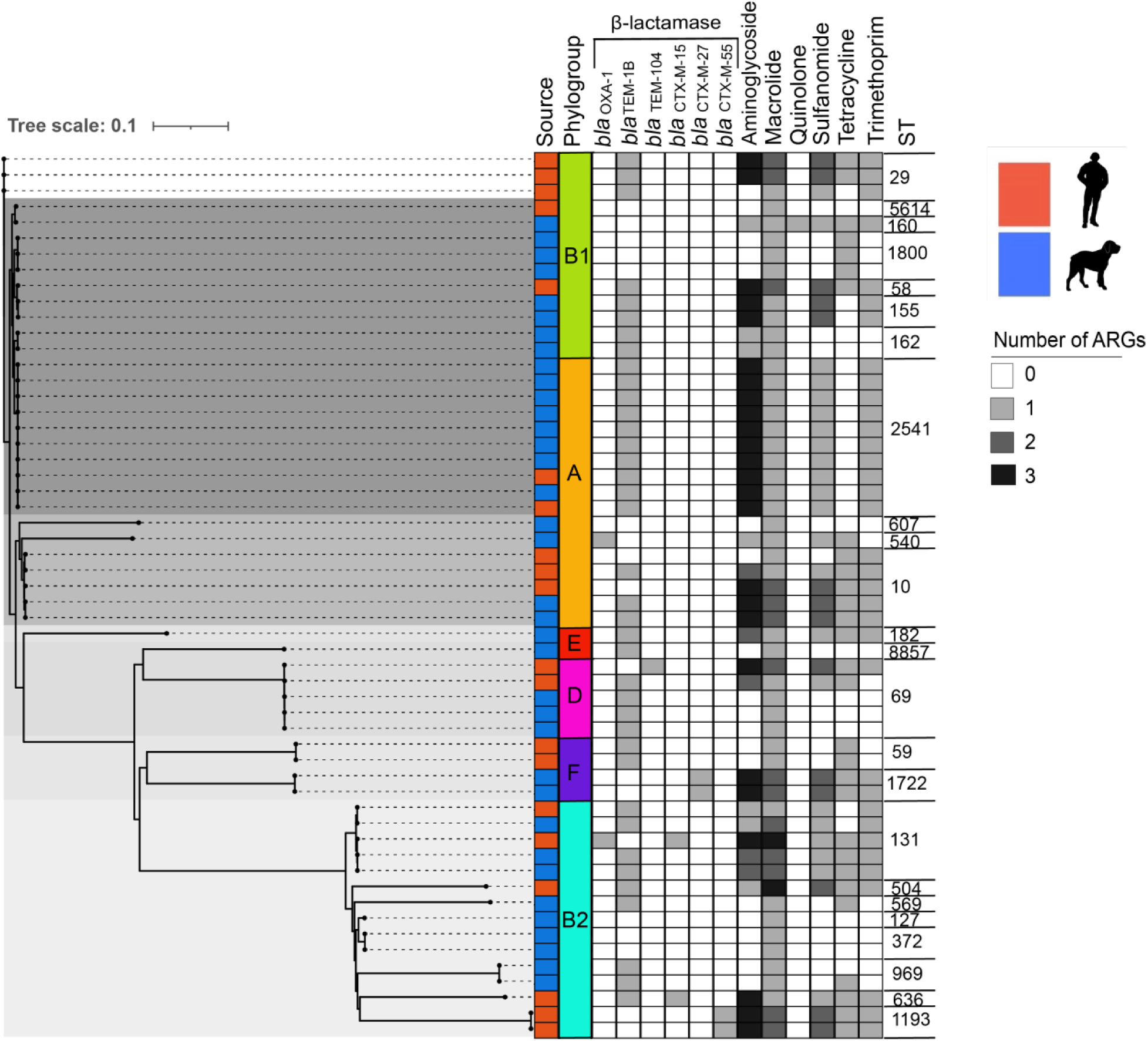
Maximum likelihood phylogenetic analysis. Phylogram depicting the best estimate of the phylogenetic relationships was computed with RAxML using 200473 SNP sites among the core genome of *E. coli* isolates with bootstrapping of 100 replicates. ClonalFrameML was used to correct the branch lengths of the tree to account for recombination. Fecal sample source, phylogroups, sequence type and ARG carriage of each isolate is indicated. Tree clades are indicated by gray shading.

Phylogroups B2 contained human isolates carrying the ESBL genes *bla*_CTX-M-55_ and *bla*_CTX-M-15_ and isolates belonging to the aforementioned pandemic lineages. Canine isolates containing ESBL gene *bla*_CTX-M-27_ belonged to phylogroup F.

Core genome MLST revealed six epidemiologically linked clonal groups using previously defined thresholds. All clonal groups contained isolates carrying the β-lactamase gene, *bla*_TEM-1B._ Five of the sharing groups contained two isolate pairs that originated from canine fecal samples. Three of the sharing groups belonged to the pandemic lineages ST131, ST69 and ST10. Sharing group 2 belonging to ST69 contained isolates sampled on different days. The largest clonal group was sharing group 6 with isolates originating from both human (n=1) and canine (n=5) fecal samples and spanning three sampling dates.

By performing Bayesian phylogenetic inference from local outbreak clusters using MASCOT, we found support for at least two host jumps of *E. coli* between humans and canines (Figure 5A). In particular, one of the local outbreak clusters shows that human samples are nested in a clade of canine sequences (Figure 5B). Additionally, we find a second outbreak cluster with sequences from humans and canines, which indicates a second host jump. In this case, however, the root of the local outbreak clusters lies several years in the past, and as such, may have taken different routes of transmission that are not accounted for in the model. The different *E. coli* isolates contained largely consistent plasmid profiles within the same local outbreak clusters (Figure 5C).

**Figure 5.**
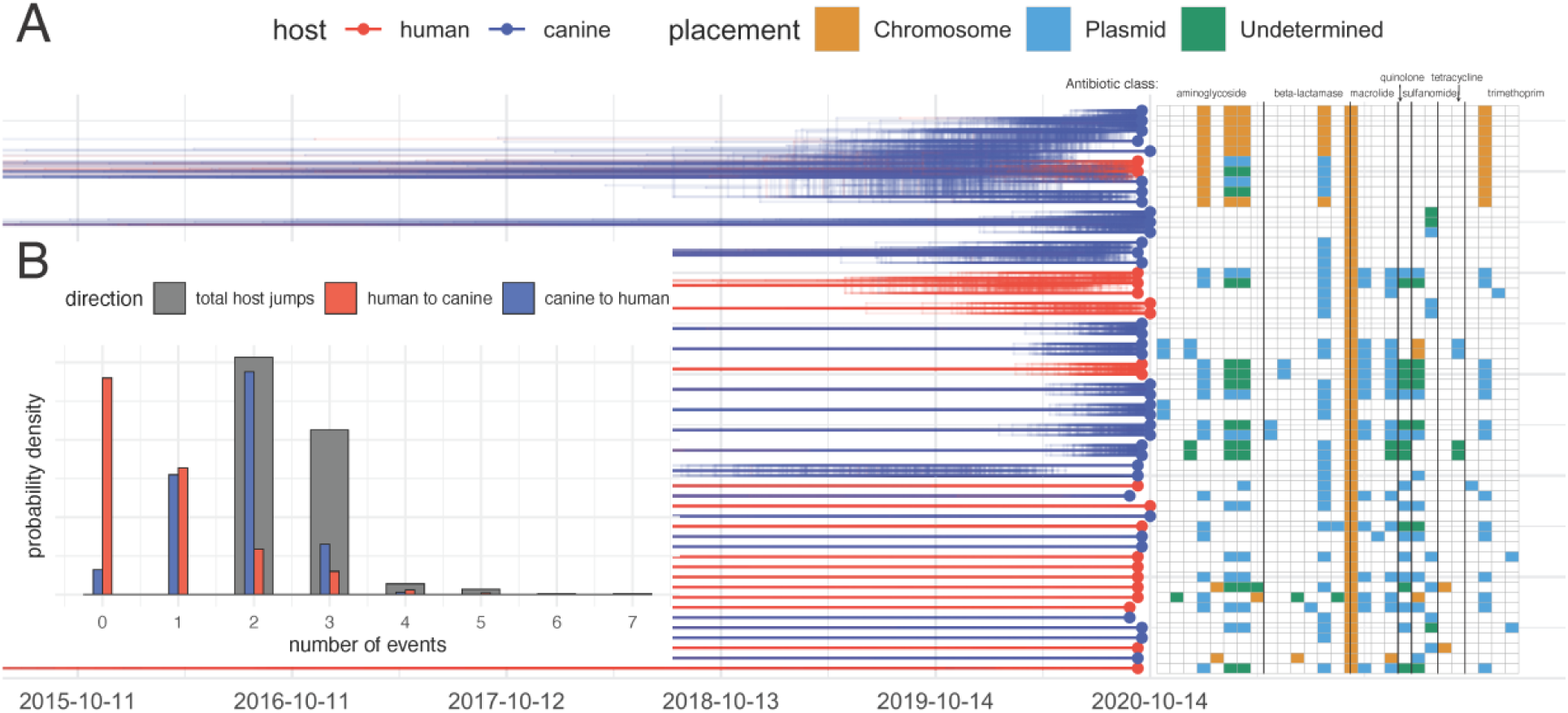
Transmission of *E. coli* between canine and humans. (A) Inferred number of host jumps. Host jumps are computed as the number of edges for which parent and child nodes are in different states. The total number denotes the posterior distribution of host jump events in either direction. (B) Posterior distribution of typed phylogenetic trees inferred from local outbreak clusters using MASCOT. The trees show the densitree representation of the local outbreak clusters plotted using ggtree. Resistance gene profile for isolates are depicted on the right, with orange tiles representing a resistance gene to be chromosomally located, blue tiles representing a resistance gene to be plasmid located and green tiles representing an undetermined resistance gene location.

## Discussion

There remains a gap in the literature regarding the role of canines in environmental contamination in urban settings of high-income countries. Therefore, we aimed to investigate antibiotic resistance of canine feces on San Francisco streets and its relatedness to human fecal samples in the same environment. Our results demonstrated that on average humans presented a 0.5 times higher proportion of antibiotic resistant *E. coli* (ABR-Ec) isolates compared to canines, but the ARG repertoire of the ABR-Ec isolates were similar between the two species. We were surprised to find that despite the supposedly high reports of human fecal waste in San Francisco (54), the majority of our samples belonged to canines.

Our results show that the risk of ABR enteric pathogen carriage was similar between species, in accordance with overall pathogen carriage from previous research (23). Aside from one isolate in canines determined to be diffuse adherent *E. coli,* neither species carried any of the six diarrheagenic *E. coli* pathotypes based on previously defined definitions. A comparable percentage of *E. coli* in both species did carry the virulence gene *eae,* which is known to enhance virulence of STEC infections (55), but did not carry the additional *stx1/stx2* or *bfp* genes of STEC and typical enteropathogenic *E. coli*, respectively. Both human and canine *E. coli* showed high rates of ampicillin and trimethoprim-sulfamethoxazole phenotypic resistance, suggesting the need for drug monitoring and appropriate administration in the community. The lack of cefazolin specific resistance genes in both species was surprising considering the high rate of phenotypic resistance to the drug. Cefazolin resistance has been shown to be mediated by *ampC* β-lactamases (56), which were not detected in this study group. The high counts of *bla*_TEM-1B_, known to confer resistance to structurally similar first-generation cephalosporin, cephalothin (57), in addition to the presence of other β-lactamase genes are most likely are conferring resistance to cefazolin.

In alignment with previous studies (58), (14), (12) our results also show there was a high degree of similarity between the resistance genes and plasmids found in human and canine samples. The most frequent ARG in both species was the broad-spectrum macrolide gene *mdf(A)*, followed by the non-ESBL gene, *bla*_TEM-1B_. Only 50% of the ARGs were present in both species but their presence was not significantly different. The limited sample size may explain why some resistance genes were only detected in one species and why no significant difference in gene presence was found. Nevertheless, the high degree of similarity in the distribution and prevalence of shared ARGs suggests a similar composition in the gut microbiome of the two species despite recent findings challenging this notion (16). This discrepancy may be explained by multiple factors such as environmental setting and sampling window, as *E. coli* presence in the gut is known to be highly dynamic due to selection pressure and clonal competition (59).

Both species had a high prevalence of the Col156 and IncFIB plasmids in addition to various species-specific plasmids. The ESBL genes, *bla*_CTX-M-55_ and *bla*_CTX-M-27_ were exclusively found on plasmid-called contigs in humans and canines, respectively. The missing replicon assignment to the majority of ARG carrying plasmid fragments is most likely due to a lack of plasmid markers on the short-read contigs (60). Nevertheless, three IncF-type plasmids harboring the non-ESBL *bla*_TEM-1B_ gene were found in both species. IncF plasmids are well known to disseminate a range of resistance genes including *bla*_CTX-M-15_ (61) while being stably maintained without antibiotic pressure (62). This suggests plasmid mediated ARG exchange and persistence between the two species may be occurring regardless of antibiotic usage in the community.

This potential for ARG exchange is further supported by the overlap in clonal species between the two sources. The maximum-likelihood phylogenetic tree showed no clustering by species, with canine and human isolates equally distributed across sequence types and phylogroups. Core genome MLST found five epidemiologically related clonal groups across canine fecal samples, three of which belonged to pandemic linages ST131, ST69, and ST10. The clonal group ST2541 contained epidemiologically linked isolates from both human and canines over multiple sample dates, suggesting recent clonal dissemination across the two species. Past studies in Brazil (63) and Norway (64) have found ST2541 to be prevalent in stray and domestic canines, but to our knowledge this is the first instance of it occurring in both humans and canines of the same study base.

Our phylodynamic analyses show that we can utilize genomic sequencing to reconstruct host jump events on a local scale. Our analyses also suggest that there is cross-species transmission of *E. coli* between canines and humans, with the samples in the same outbreak clusters having the same or very similar plasmid profiles. Similar plasmid profiles between local outbreak clusters inferred from chromosomal DNA suggest that at the time scales we are investigating, plasmids are largely maintained even during cross-species transmission. Using more isolates or explicitly tracking the movement of plasmids between different bacterial lineages could shine additional light into the cross-species transmission of *E. coli* (44). With relatively few host jumps captured by the genomic data, quantifying the rates of host jumps is complex. This could be enabled by larger datasets and could be used to, for example, estimate the number of cases in each species directly caused by host jumps (41), (45). Knowing these rates could potentially be used to parameterize epidemiological models to predict the impact of interventions to reduce the burden of ARB gene carrying *E coli*.

One possible explanation for the high similarity and transmission of the ABR-*E. coli* in humans and canines is a cohabitation between the two species such as canine ownership, which has been shown to contribute to *E. coli* sharing and long-term colonization (66), (64), (67). Alternatively, it is also possible the fecal samples originated from canines unrelated to the human samples such as passersby’s walking their canines or strays. It is also possible the non-human fecal samples originated from animals other than canines, though this is less likely due to the limited wildlife ecology of downtown San Francisco and the appearance of the feces. The human fecal samples most likely originated from open defecation due to lack of access to sanitation facilities for the unhoused population in San Francisco (54). Individuals experiencing homelessness have been shown to be at greater risk for exposure to infectious diseases due to compounding factors (68), (69). Residence in areas where fecal contamination is prevalent, such as sidewalks, can lead to increased environmental exposure to ABR-*E. coli* resulting in spillover, gene exchange and *E. coli* colonization from canine feces.

A high number of canines in San Francisco has been highlighted and this highlights the need for public health interventions to minimize potential environmental contamination. A limitation of this study was that we were not able to gather individual data for fecal samples, so it is possible some fecal samples originated from the same human or canine overtime. We were also not able to establish the extent of contact between the individuals contributing to the fecal samples, which would help support transmission events and the role of the environment as a medium between the two species.

In conclusion, our study found a high degree of similarity in ABR-*E. coli* in human and canine fecal samples on the sidewalks of San Francisco as well as recent transmission events from canines to humans. Our results support the wider use of phylodynamic methods in bacterial surveillance to refine insights on ARB contribution and highlight routes of transmission that may warrant intervention. We discovered a wide variety of resistance genes including a high prevalence of macrolide and β-lactamase genes in canines. The high degree of overlap between the human and canine phenotypic and genotypic resistance suggests domesticated and stray canines play an important role as reservoir and vector for environmental contamination of ARGs. Our results also support clonal spread of ABR-*E. coli* from canines to humans in San Francisco, most likely through the environment. Canines are required to be on leash when off personal property, and owners are required to remove and dispose properly of any feces. Despite these ordinances there remains a high frequency of canine feces on San Francisco sidewalks. This study supports public health efforts to report and remove both human and canine feces on city streets, such as San Francisco’s 311 program, as well as increased signage and enforcement of ordinances. Public health measures should be continued in order to reduce ARB spillover to the environment and risk of exchange to humans.

## Acknowledgments

N.F.M. is supported by NIH NIGMS R35 GM119774

J.P.G. is supported on NIH NIAID R01AI167989

## Abbreviation

ABR: Antibiotic resistance
ARB: Antibiotic resistant bacteria
ABR-Ec: Antibiotic resistant *Escherichia coli*
ARG: Antibiotic resistance genes
ESBL: Extended spectrum β-lactamases
MDR: Multidrug resistant

## Notes

### Competing Interest Statement

The authors have declared no competing interest.

## References

1. Centers for Disease Control and Prevention (U.S.). Antibiotic resistance threats in the United States, 2019 [Internet]. Centers for Disease Control and Prevention (U.S.); 2019 Nov [cited 2022 Dec 30]. Available from: https://stacks.cdc.gov/view/cdc/82532

2. Murray CJ, Ikuta KS, Sharara F, Swetschinski L, Robles Aguilar G, Gray A, et al. Global burden of bacterial antimicrobial resistance in 2019: a systematic analysis. The Lancet. 2022 Feb;399(10325):629–55.

3. Woolhouse M, Ward M, van Bunnik B, Farrar J. Antimicrobial resistance in humans, livestock and the wider environment. Phil Trans R Soc B. 2015 Jun 5;370(1670):20140083.

4. AbuOun M, O’Connor HM, Stubberfield EJ, Nunez-Garcia J, Sayers E, Crook DW, et al. Characterizing Antimicrobial Resistant Escherichia coli and Associated Risk Factors in a Cross-Sectional Study of Pig Farms in Great Britain. Front Microbiol. 2020 May 25;11:861.

5. Tenaillon O, Skurnik D, Picard B, Denamur E. The population genetics of commensal Escherichia coli. Nat Rev Microbiol. 2010 Mar;8(3):207–17.

6. Alvarez-Uria G, Gandra S, Mandal S, Laxminarayan R. Global forecast of antimicrobial resistance in invasive isolates of Escherichia coli and Klebsiella pneumoniae. International Journal of Infectious Diseases. 2018 Mar;68:50–3.

7. Carattoli A. Resistance Plasmid Families in Enterobacteriaceae. Antimicrob Agents Chemother. 2009 Jun;53(6):2227–38.

8. Zhang S, Abbas M, Rehman MU, Huang Y, Zhou R, Gong S, et al. Dissemination of antibiotic resistance genes (ARGs) via integrons in Escherichia coli: A risk to human health. Environmental Pollution. 2020 Nov;266:115260.

9. Robinson TP, Bu DP, Carrique-Mas J, Fèvre EM, Gilbert M, Grace D, et al. Antibiotic resistance is the quintessential One Health issue. Trans R Soc Trop Med Hyg. 2016 Jul;110(7):377–80.

10. Musoke D, Namata C, Lubega GB, Niyongabo F, Gonza J, Chidziwisano K, et al. The role of Environmental Health in preventing antimicrobial resistance in low-and middle-income countries. Environ Health Prev Med. 2021 Dec;26(1):100.

11. Reinthaler FF, Posch J, Feierl G, Wüst G, Haas D, Ruckenbauer G, et al. Antibiotic resistance of E. coli in sewage and sludge. Water Research. 2003 Apr;37(8):1685–90.

12. Damborg P, Morsing MK, Petersen T, Bortolaia V, Guardabassi L. CTX-M-1 and CTX-M-15-producing Escherichia coli in dog faeces from public gardens. Acta Vet Scand. 2015 Dec;57(1):83.

13. Marchetti L, Buldain D, Gortari Castillo L, Buchamer A, Chirino-Trejo M, Mestorino N. Pet and Stray Dogs as Reservoirs of Antimicrobial-Resistant Escherichia coli. Prasad S, editor. International Journal of Microbiology. 2021 Jan 25;2021:1–8.

14. Zhao R, Hao J, Yang J, Tong C, Xie L, Xiao D, et al. The co-occurrence of antibiotic resistance genes between dogs and their owners in families. iMeta [Internet]. 2022 Jun [cited 2022 Jul 10];1(2). Available from: https://onlinelibrary.wiley.com/doi/10.1002/imt2.21

15. Johnson JR, Kaster N, Kuskowski MA, Ling GV. Identification of Urovirulence Traits in Escherichia coli by Comparison of Urinary and Rectal E. coli Isolates from Dogs with Urinary Tract Infection. J Clin Microbiol. 2003 Jan;41(1):337–45.

16. Røken M, Forfang K, Wasteson Y, Haaland AH, Eiken HG, Hagen SB, et al. Antimicrobial resistance—Do we share more than companionship with our dogs? J of Applied Microbiology. 2022 May 29;jam.15629.

17. Muloi D, Ward MJ, Pedersen AB, Fèvre EM, Woolhouse MEJ, van Bunnik BAD. Are Food Animals Responsible for Transfer of Antimicrobial-Resistant Escherichia coli or Their Resistance Determinants to Human Populations? A Systematic Review. Foodborne Pathogens and Disease. 2018 Aug;15(8):467–74.

18. Keller A, Ankenbrand MJ. Inferring Core Genome Phylogenies for Bacteria. In: Mengoni A, Bacci G, Fondi M, editors. Bacterial Pangenomics [Internet]. New York, NY: Springer US; 2021 [cited 2023 May 2]. p. 59–68. (Methods in Molecular Biology; vol. 2242). Available from: https://link.springer.com/10.1007/978-1-0716-1099-2_4

19. Ingle DJ, Howden BP, Duchene S. Development of Phylodynamic Methods for Bacterial Pathogens. Trends in Microbiology. 2021 Sep;29(9):788–97.

20. Guinat C, Vergne T, Kocher A, Chakraborty D, Paul MC, Ducatez M, et al. What can phylodynamics bring to animal health research? Trends in Ecology & Evolution. 2021 Sep;36(9):837–47.

21. Pokharel S, Raut S, Adhikari B. Tackling antimicrobial resistance in low-income and middle-income countries. BMJ Glob Health. 2019 Nov;4(6):e002104.

22. Knee J, Sumner T, Adriano Z, Anderson C, Bush F, Capone D, et al. Effects of an urban sanitation intervention on childhood enteric infection and diarrhea in Maputo, Mozambique: A controlled before-and-after trial. eLife. 2021 Apr 9;10:e62278.

23. Barker T, Capone D, Amato HK, Clark R, Henderson A, Holcomb DA, et al. Public toilets have reduced enteric pathogen hazards in San Francisco [Internet]. Public and Global Health; 2023 Feb [cited 2023 Mar 13]. Available from: http://medrxiv.org/lookup/doi/10.1101/2023.02.10.23285757

24. CLSI M100-Performance Standards for Antimicrobial Suceptibility Testing.

25. Wick RR, Judd LM, Gorrie CL, Holt KE. Unicycler: Resolving bacterial genome assemblies from short and long sequencing reads. Phillippy AM, editor. PLoS Comput Biol. 2017 Jun 8;13(6):e1005595.

26. Mikheenko A, Prjibelski A, Saveliev V, Antipov D, Gurevich A. Versatile genome assembly evaluation with QUAST-LG. Bioinformatics. 2018 Jul 1;34(13):i142–50.

27. Seemann T. Abricate [Internet]. Github; Available from: https://github.com/tseemann/abricate

28. Vidal M, Kruger E, Durán C, Lagos R, Levine M, Prado V, et al. Single Multiplex PCR Assay To Identify Simultaneously the Six Categories of Diarrheagenic Escherichia coli Associated with Enteric Infections. J Clin Microbiol. 2005 Oct;43(10):5362–5.

29. Arredondo-Alonso S, Rogers MRC, Braat JC, Verschuuren TD, Top J, Corander J, et al. mlplasmids: a user-friendly tool to predict plasmid-and chromosome-derived sequences for single species. Microbial Genomics [Internet]. 2018 Nov 1 [cited 2021 Sep 3];4(11). Available from: https://www.microbiologyresearch.org/content/journal/mgen/10.1099/mgen.0.000224

30. Robertson J, Nash JHE. MOB-suite: software tools for clustering, reconstruction and typing of plasmids from draft assemblies. Microbial Genomics [Internet]. 2018 Aug 1 [cited 2022 May 11];4(8). Available from: https://www.microbiologyresearch.org/content/journal/mgen/10.1099/mgen.0.000206

31. Seemann T. mlst [Internet]. Github; Available from: https://github.com/tseemann/mlst

32. Jolley KA, Maiden MC. BIGSdb: Scalable analysis of bacterial genome variation at the population level. BMC Bioinformatics. 2010 Dec;11(1):595.

33. Clausen PTLC, Aarestrup FM, Lund O. Rapid and precise alignment of raw reads against redundant databases with KMA. BMC Bioinformatics. 2018 Dec;19(1):307.

34. Kluytmans-van den Bergh MFQ, Rossen JWA, Bruijning-Verhagen PCJ, Bonten MJM, Friedrich AW, Vandenbroucke-Grauls CMJE, et al. Whole-Genome Multilocus Sequence Typing of Extended-Spectrum-Beta-Lactamase-Producing Enterobacteriaceae. Ledeboer NA, editor. J Clin Microbiol. 2016 Dec;54(12):2919–27.

35. Waters, Nick, Pritchard, Leighton. EzClermont [Internet]. Available from: https://github.com/nickp60/EzClermont

36. Seemann T. Prokka: rapid prokaryotic genome annotation. Bioinformatics. 2014 Jul 15;30(14):2068–9.

37. Page AJ, Cummins CA, Hunt M, Wong VK, Reuter S, Holden MTG, et al. Roary: rapid large-scale prokaryote pan genome analysis. Bioinformatics. 2015 Nov 15;31(22):3691–3.

38. Page AJ, Taylor B, Delaney AJ, Soares J, Seemann T, Keane JA, et al. SNP-sites: rapid efficient extraction of SNPs from multi-FASTA alignments. Microbial Genomics [Internet]. 2016 Apr 29 [cited 2022 Jul 14];2(4). Available from: https://www.microbiologyresearch.org/content/journal/mgen/10.1099/mgen.0.000056

39. Stamatakis A. RAxML version 8: a tool for phylogenetic analysis and post-analysis of large phylogenies. Bioinformatics. 2014 May 1;30(9):1312–3.

40. Didelot X, Wilson DJ. ClonalFrameML: Efficient Inference of Recombination in Whole Bacterial Genomes. Prlic A, editor. PLoS Comput Biol. 2015 Feb 12;11(2):e1004041.

41. Letunic I, Bork P. Interactive Tree Of Life (iTOL): an online tool for phylogenetic tree display and annotation. Bioinformatics. 2007 Jan 1;23(1):127–8.

42. Müller NF, Wagner C, Frazar CD, Roychoudhury P, Lee J, Moncla LH, et al. Viral genomes reveal patterns of the SARS-CoV-2 outbreak in Washington State. Sci Transl Med. 2021 May 26;13(595):eabf0202.

43. Müller NF, Wüthrich D, Goldman N, Sailer N, Saalfrank C, Brunner M, et al. Characterising the epidemic spread of influenza A/H3N2 within a city through phylogenetics. Lauring AS, editor. PLoS Pathog. 2020 Nov 19;16(11):e1008984.

44. Müller NF, Duchêne S, Williamson DA, Bedford T, Howden BP, Ingle DJ. Tracking the horizontal transfer of plasmids in Shigella sonnei and Shigella flexneri using phylogenetics [Internet]. Evolutionary Biology; 2022 Oct [cited 2023 May 3]. Available from: http://biorxiv.org/lookup/doi/10.1101/2022.10.27.514108

45. Bouckaert RR. DensiTree: making sense of sets of phylogenetic trees. Bioinformatics. 2010 May 15;26(10):1372–3.

46. Yu G, Smith DK, Zhu H, Guan Y, Lam TT. GGTREE : an R package for visualization and annotation of phylogenetic trees with their covariates and other associated data. McInerny G, editor. Methods Ecol Evol. 2017 Jan;8(1):28–36.

47. Wickham H. ggplot2: Elegant Graphics for Data Analysis [Internet]. Springer-Verlag New York; 2016. Available from: https://ggplot2.tidyverse.org

48. Wickham H, Francois R, Henry L, Müller K. dplyr: A Grammar of Data Manipulation [Internet]. Vol. R package version 1.0.3. 2021. Available from: https://CRAN.R-project.org/package=dplyr

49. Wickham H. stringr: Simple Consistent Wrappers for Common String Operations [Internet]. Vol. R package version 1.4.0. 2019. Available from: https://CRAN.R-project.org/package=stringr

50. Wickham H. tidyr: Tidy Messy Data [Internet]. Vol. R package version 1.1.2. 2020. Available from: https://CRAN.R-project.org/package=tidyr

51. Sjoberg, David. ggsankey: Sankey, Alluvial and Sankey Bump Plots. 2022.

52. QGIS Association. QGIS Geographic Information System [Internet]. 2022. Available from: http://www.qgis.org

53. Zhu K, Suttner B, Pickering A, Konstantinidis KT, Brown J. A novel droplet digital PCR human mtDNA assay for fecal source tracking. Water Research. 2020 Sep;183:116085.

54. Amato HK, Martin D, Hoover CM, Graham JP. Somewhere to go: assessing the impact of public restroom interventions on reports of open defecation in San Francisco, California from 2014 to 2020. BMC Public Health. 2022 Sep 4;22(1):1673.

55. Hua Y, Bai X, Zhang J, Jernberg C, Chromek M, Hansson S, et al. Molecular characteristics of eae-positive clinical Shiga toxin-producing Escherichia coli in Sweden. Emerging Microbes & Infections. 2020 Jan 1;9(1):2562–70.

56. Kawamura M, Ito R, Tamura Y, Takahashi M, Umenai M, Chiba Y, et al. Overproduction of Chromosomal ampC β-Lactamase Gene Maintains Resistance to Cefazolin in Escherichia coli Isolates. Wang H, editor. Microbiol Spectr. 2022 Jun 29;10(3):e00058–22.

57. Wick WE, Preston DA. Biological Properties of Three 3-Heterocyclic-Thiomethyl Cephalosporin Antibiotics. Antimicrob Agents Chemother. 1972 Mar;1(3):221–34.

58. Salinas L, Loayza F, Cárdenas P, Saraiva C, Johnson TJ, Amato H, et al. Environmental Spread of Extended Spectrum Beta-Lactamase (ESBL) Producing Escherichia coli and ESBL Genes among Children and Domestic Animals in Ecuador. Environ Health Perspect. 2021 Feb;129(2):027007.

59. Loayza F, Graham JP, Trueba G. Factors Obscuring the Role of E. coli from Domestic Animals in the Global Antimicrobial Resistance Crisis: An Evidence-Based Review. IJERPH. 2020 Apr 28;17(9):3061.

60. Juraschek K, Borowiak M, Tausch SH, Malorny B, Käsbohrer A, Otani S, et al. Outcome of Different Sequencing and Assembly Approaches on the Detection of Plasmids and Localization of Antimicrobial Resistance Genes in Commensal Escherichia coli. Microorganisms. 2021 Mar 14;9(3):598.

61. Kim J, Bae IK, Jeong SH, Chang CL, Lee CH, Lee K. Characterization of IncF plasmids carrying the blaCTX-M-14 gene in clinical isolates of Escherichia coli from Korea. Journal of Antimicrobial Chemotherapy. 2011 Jun 1;66(6):1263–8.

62. Lucas P, Jouy E, Le Devendec L, de Boisséson C, Perrin-Guyomard A, Jové T, et al. Characterization of plasmids harboring blaCTX-M genes in Escherichia coli from French pigs. Veterinary Microbiology. 2018 Oct;224:100–6.

63. Melo LC, Oresco C, Leigue L, Netto HM, Melville PA, Benites NR, et al. Prevalence and molecular features of ESBL/pAmpC-producing Enterobacteriaceae in healthy and diseased companion animals in Brazil. Veterinary Microbiology. 2018 Jul;221:59–66.

64. Toombs-Ruane LJ, Benschop J, French NP, Biggs PJ, Midwinter AC, Marshall JC, et al. Carriage of Extended-Spectrum-Beta-Lactamase-and AmpC Beta-Lactamase-Producing Escherichia coli Strains from Humans and Pets in the Same Households. Dozois CM, editor. Appl Environ Microbiol. 2020 Nov 24;86(24):e01613–20.

65. Paredes MI, Perofsky AC, Frisbie L, Moncla LH, Roychoudhury P, Xie H, et al. Local-Scale phylodynamics reveal differential community impact of SARS-CoV-2 in metropolitan US county [Internet]. Infectious Diseases (except HIV/AIDS); 2022 Dec [cited 2023 May 3]. Available from: http://medrxiv.org/lookup/doi/10.1101/2022.12.15.22283536

66. Johnson JR, Davis G, Clabots C, Johnston BD, Porter S, DebRoy C, et al. Household Clustering of Escherichia coli Sequence Type 131 Clinical and Fecal Isolates According to Whole Genome Sequence Analysis. Open Forum Infectious Diseases. 2016 May 1;3(3):ofw129.

67. Habib I, Mohteshamuddin K, Mohamed MYI, Lakshmi GB, Abdalla A, Bakhit Ali Alkaabi A. Domestic Pets in the United Arab Emirates as Reservoirs for Antibiotic-Resistant Bacteria: A Comprehensive Analysis of Extended-Spectrum Beta-Lactamase Producing Escherichia coli Prevalence and Risk Factors. Animals. 2023 May 9;13(10):1587.

68. Fazel S, Geddes JR, Kushel M. The health of homeless people in high-income countries: descriptive epidemiology, health consequences, and clinical and policy recommendations. The Lancet. 2014 Oct;384(9953):1529–40.

69. Liu CY, Chai SJ, Watt JP. Communicable disease among people experiencing homelessness in California. Epidemiol Infect. 2020;148:e85.

