## Supplemental tables for "Phylodynamics Uncovers the Transmission of Antibiotic-Resistant *Escherichia coli* between Canines and Humans in an Urban Environment"

### Supplemental Table 1: Distribution of Plasmid Replicons in Human and Canine Isolates

| Replicon Group | Plasmid Replicon Type | Percent of Human Isolates (%) | Percent of Canine Isolates (%) |
|---|---|---|---|
| Col | Col(BS512) | 30.0 | 0.0 |
| | Col(MG828) | 40.0 | 5.6 |
| | Col156 | 75.0 | 41.7 |
| | Col440I | 5.0 | 0.0 |
| | Col440II | 5.0 | 0.0 |
| | Col8282 | 30.0 | 30.6 |
| | ColpVC | 15.0 | 0.0 |
| | ColRNAI | 70.0 | 36.1 |
| IncB | IncB/O/K/Z | 30.0 | 0.0 |
| IncF | IncFIA | 25.0 | 13.9 |
| | IncFIB(AP001918) | 65.0 | 55.6 |
| | IncFIC(FII) | 35.0 | 11.1 |
| | IncFII | 0.0 | 16.7 |
| | IncFII(29) | 30.0 | 25.0 |
| | IncFII(pCoo) | 15.0 | 8.3 |
| | IncFII(pCRY) | 5.0 | 0.0 |
| | IncFII(pHN7A8) | 5.0 | 0.0 |
| | IncFII(pRSB107) | 10.0 | 8.3 |
| IncI | IncI1_1_Alpha | 50.0 | 11.1 |
| | IncI2_1_Delta | 10.0 | 0.0 |
| | IncL/M(pMU407) | 5.0 | 0.0 |
| IncX | IncX1 | 0.0 | 2.8 |
| | IncX4 | 20.0 | 0.0 |
| IncY | IncY | 0.0 | 8.3 |
| Other | p0111 | 0.0 | 11.1 |
| | pENTAS02 | 0.0 | 2.8 |
| | RepA_1_pKPC-CAV1321 | 0.0 | 2.8 |

### Supplemental Table 2: Diarrheagenic Virulence Gene Carriage in Human and Canina Isolates

| Virulence Gene | ST (n) | Percent of Human Isolates (n=20) | Percent of Canine Isolates (n=36) | P-value* |
|---|---|---|---|---|
| *eae* | 10 (2), 29 (3), 2541 (10) | 25 | 27 | 1.0 |
| *daaE* | 131 (1) | 0 | 2.7 | 1.0 |
| *alpha of 0.05 | | | | |
